# Involvement of sensory neuron-TRPV4 in acute and chronic itch behaviors

**DOI:** 10.1101/2021.07.06.451366

**Authors:** Quan Fang, Zilong Wang, Abbie Suttle, Yong Chen

**Affiliations:** Departments of Neurology, Duke University, Durham, NC 27710, USA; Anesthesiology, Duke University, Durham, NC 27710, USA; Pathology, Duke University, Durham, NC 27710, USA

**Author notes:** Correspondence author: Yong Chen), Department of Neurology, Duke University, Durham, NC 27710, USA.

## Abstract

Itch, particularly chronic itch, negatively impacts patients’ physical, social, and psychological well-being, leading to deterioration in their quality of life. Limited understanding of itch mechanisms hinders the development of effective antipruritic treatments.TRPV4, a multimodally activated nonselective cation channel, has been detected in sensory neurons of dorsal root and trigeminal ganglion (DRG, TG) and skin cells (e.g. keratinocytes, mast cells, and macrophages). Recent evidence from experimental and clinical relevant studies has implicated that TRPV4 in skin cells plays an important role in both acute and chronic itch. In contrast, little is known whether TRPV4 in sensory neurons directly contributes to scratching behaviors. Here we used sensory neuron-Trpv4 conditional knockout (cKO) mice to address this question. Our results showed that TRPV4 in sensory neurons contributes to scratching behavior evoked by histaminergic (histamine and 48/80) and partial histaminergic (5-HT), but not non-histaminergic (SLIGRL and CQ) pruritogens. Moreover, we observed that TRPV4 in sensory neurons is required for dry skin, but not allergic contact dermatitis, -associated chronic itch. These findings suggest that neuronal-TRPV4 might be specific for some forms of acute and chronic itch.

## INTRODUCTION

Limited understanding of itch mechanisms hinders development of antipruritic treatments. Accumulating evidence suggests that transient receptor potential (TRP) ion channels as potential therapeutic targets for itch (1–3). Relative to the extensive studies on TRPV1 and TRPA in itch, TRPV4 has received little research attention until recently. TRPV4, a multimodally activated nonselective cation channel, has been detected in sensory neurons of dorsal root and trigeminal ganglion (DRG, TG) and skin cells (4–6), which are involved in itch sensation and development. Using *Trpv4* knockout (KO) mice, several groups reported that it is required for pruritogens-evoked acute itch (7–9). To determine cellular sites of action of *Trpv4* in itch, we employed keratinocyte-*Trpv4* conditional KO (cKO) mice and found that deletion of *Trpv4* significantly reduced histaminergic (histamine and 48/80), but not non-histaminergic (chloroquine, CQ) pruritogens, -evoked acute itch (8). Additionally, we and others have recently revealed that TRPV4 in skin keratinocytes and macrophages also critically mediates chronic itch (4, 5). Although these studies strongly suggest that TRPV4 in skin cells is essential for itch, whether neuronal-TRPV4 directly contributes to acute and chronic itch behaviors remains largely elusive. Here we were prompted to use sensory neuron-*Trpv4* cKOs to address this question.

## MATERIALS AND METHODS

Mice with conditional deletion for *Trpv4* in primary sensory neurons were generated by mating *Trpv4^fl/fl^* with *Nav1.8-Cre* mice, as described previously (4). Efficiency of targeting *Trpv4* was verified in both DRGs and TGs at gene and protein levels, see supplement data in study (4). All animal protocols were approved by the Duke University Institutional Animal Care and Use Committee (IACUC). Male *Trpv4* cKO (Nav1.8-cre::*Trpv4*^fl/fl^, C57bl/6 background) and wild-type (WT) mice were used.

All chemicals were purchased from Sigma-Aldrich. For acute itch, mice received intradermal (i.d.) injection of 50μl of pruritogens in normal saline: histamine (500μg), 48/80 (100μg), 5-HT (25μg), SLIGRL (100μg), and chloroquine (200μg), into the nape of neck. For chronic itch, dry skin model was induced by painting the neck-back skin with acetone and diethyl-ether (1:1) following water (AEW) twice daily (10). An allergic contact dermatitis model was established by applying 1-fluoro-2, 4-dinitrobenzene (DNFB) onto the neck-back skin as previously described (11). Scratch bouts were analyzed as previously described (4) in a blinded way.

## RESULTS

For acute itch, we found that cKO of *Trpv4* in sensory neurons significantly attenuated the scratching responses evoked by histamine, 48/40 and 5-HT but not SLIGRL and CQ (Fig.1).

**Fig. 1.**
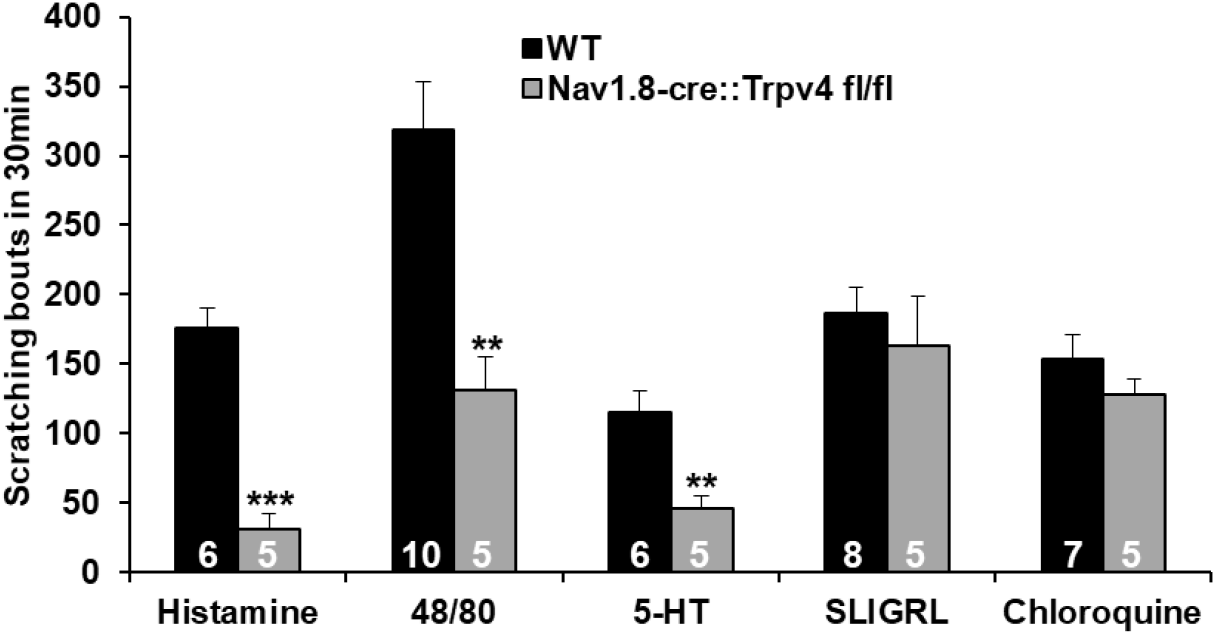
Effect of sensory neuron-TRPV4 on acute itch induced by pruritogens. Histamine, compound 48/80, and 5-HT, but not SLIGRL or chloroquine, evoked acute scratching behaviors that were significantly attenuated in *Trpv4* cKOs *vs* WTs. **p<0.01 and ***p<0.001, two-tail *t* test. Data are expressed as mean±SEM and group size is indicated in the bars.

In dry skin chronic itch model, we found that scratching behaviors were significantly reduced in *Trpv4* cKOs on days 7 and 9 after AEW treatment (Fig.2). In contrast, in allergic contact dermatitis chronic itch model, *Trpv4* cKOs did not display a significant reduction of scratching on days 12 and 14 after DNFB treatment (Fig.2).

**Fig. 2.**
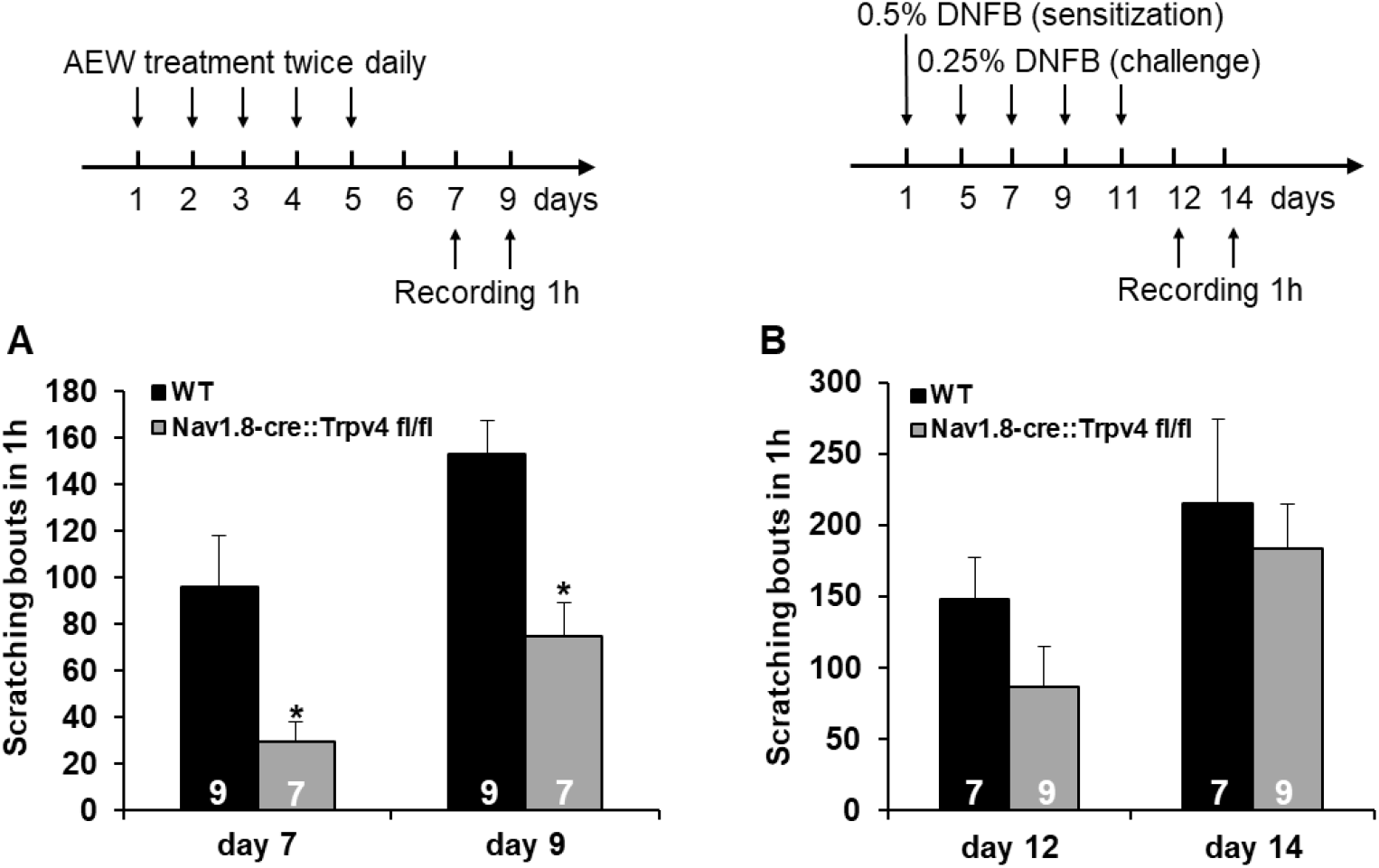
Effect of sensory neuron-TRPV4 on dry skin- or allergic contact dermatitis-associated chronic itch. Dry skin (A, AEW model), but not allergic contact dermatitis (B, DNFB model), evoked scratching behaviors that were significantly attenuated in *Trpv4* cKOs *vs* WTs. *p<0.05, two-way ANOVA with Tukey’s post-hoc test. Data are expressed as mean±SEM and group size is indicated in the bars.

## DISCUSSION

Here we determined the specific contribution of neuronal-TRPV4 to acute and chronic itch behaviors in mice. Our results showed that TRPV4 in sensory neurons contributes to scratching behavior evoked by histaminergic (histamine and 48/80) and partial histaminergic (5-HT), but not non-histaminergic (SLIGRL and CQ) pruritogens. Moreover, we observed that TRPV4 in sensory neurons is required for dry skin, but not allergic contact dermatitis, -associated chronic itch.

Several studies demonstrated that TRPV4 is involved in pruritogens-evoked acute itch, although the results across laboratories are inconsistent. Akiyama et al. reported that *Trpv4* KOs displays a reduced scratching behavior in response to 5-HT, but not to histamine and SLIGRL (PAR-2 agonist). Interestingly, they found that CQ-induced itch is even increased in *Trpv4* KOs (7). In contrast, Kim et al. showed that both histamine- and CQ-induced itch is attenuated in mice lacking *Trpv4* (9). Our group found that histamine-induced itch is decreased but CQ-induced itch remained unchanged in *Trpv4* KOs (8). Although these inconsistent findings imply TRPV4 plays a role in regulating acute itch, the crucial cellular sites of action TRPV4 remains obscure. Here we showed that selective deletion of *Trpv4* in sensory neurons significantly attenuated histamine-induced scratching behavior, which is in line with Kim et al. (9) and our (8) studies but in contrast to Akiyama’s finding (7) using *Trpv4* KOs. In support of Akiyama’s report, we observed that cKO of neuronal-*Trpv4* significantly reduced 5-HT, but not SLIGRL-evoked itch. Although CQ-induced itch has been reported as increased (7), decreased (9) or unchanged (8) in *Trpv4* KOs, we here showed that it is not significantly altered in sensory neuron-*Trpv4* cKOs. Our data suggests that sensory neurons in DRG/TG are an important locale where TRPV4 differentially regulates various forms of acute itch. Additionally, the differences in certain types of acute itch between *Trpv4* KO and sensory neuron-*Trpv4* cKO point to the possible contributions of TRPV4 in other cell lineages, such as skin macrophages, mast cells, and endothelial cells (2), to itch.

Using AEW-induced dry skin and DNFB-induced allergic contact dermatitis mouse models, we next elucidated the specific contribution of neuronal-TRPV4 to chronic itch. Luo et al. recently reported that both dry skin- and allergic contact dermatitis-associated itch is attenuated in *Trpv4* KOs (5). They further found that dry skin-associated itch is reduced in keratinocyte, but not macrophage-*Trpv4* cKOs and allergic contact dermatitis-associated itch is reduced in macrophage, but not keratinocyte-*Trpv4* cKOs. Here we found *Trpv4* deficiency in sensory neurons leads to a reduction in dry skin, but not contact allergic dermatitis-associated itch. Together, these results imply both skin keratinocytes and sensory neurons contribute to the dry skin-induced itch. However, allergic contact dermatitis-associated itch might rely on TRPV4 in skin macrophages but not sensory neurons.

In conclusion, we provided an initial answer for the specific contribution of neuronal-TRPV4 to acute and chronic itch behaviors. Our data suggests that TRPV4 in sensory neurons is essential for certain types of acute and chronic itch and implies different forms of itch may rely on different cellular sites of action of TRPV4.

## ACKNOWLEDGEMENTS

This work was supported by National Institutes of Health Grant R01DE027454 to YC.

## Conflicts of interest

The authors declare that they have no conflicts of interest with the contents of this article.

## Notes

### Competing Interest Statement

The authors have declared no competing interest.

